## Supplementary Results for "A Computational Framework for Understanding the Impact of Prior Experiences on Pain Perception and Neuropathic Pain"

### *Figure S1 – Study 2 from (Jepma et al., 2018) with single-layer Kalman filter*

These simulations are run as described for Figure 1 in the main manuscript, with three key changes:

1) Here, we use the experimental data from Study 2 from (Jepma et al., 2018), instead of Study 1 as in the main manuscript.
2) We have included a third, neutral cue (as in the experimental data of Study 2) by letting $\boldsymbol{u}^{(k)}$ be a three-dimensional vector such that $\boldsymbol{u}^{(k)}_{placebo} = [u\ 0\ 0]^T$, $\boldsymbol{u}^{(k)}_{nocebo} = [0\ u\ 0]^T$ and $\boldsymbol{u}^{(k)}_{neutral} = [0\ 0\ u]^T$ and $\hat{B} = [\hat{b}_{placebo}, \hat{b}_{nocebo}, \hat{b}_{neutral}\ ]$, where $\hat{b}_{neutral} \sim N(0.9, 0.8^2)$ (same as the initial value of the elements of $\hat{B}$ for the conditioning simulations for Figure 4).
3) We have altered the level of tissue damage elicited by the thermal stimulation to be $x_{low} = 2.5$ for low heat trials (47°C, $x_{low} = 3.1$ for Figure 1) and $x_{high} = 3.5$ for high heat trials (48°C, $x_{high} = 4.3$ for Figure 1), to reflect differing sensitivity to heat at different locations on the body. In Study 1, the thermal stimuli were applied to the inner forearm, and in Study 2 the site of stimulation was the lower leg (which is less sensitive, hence the lower values of $x$).

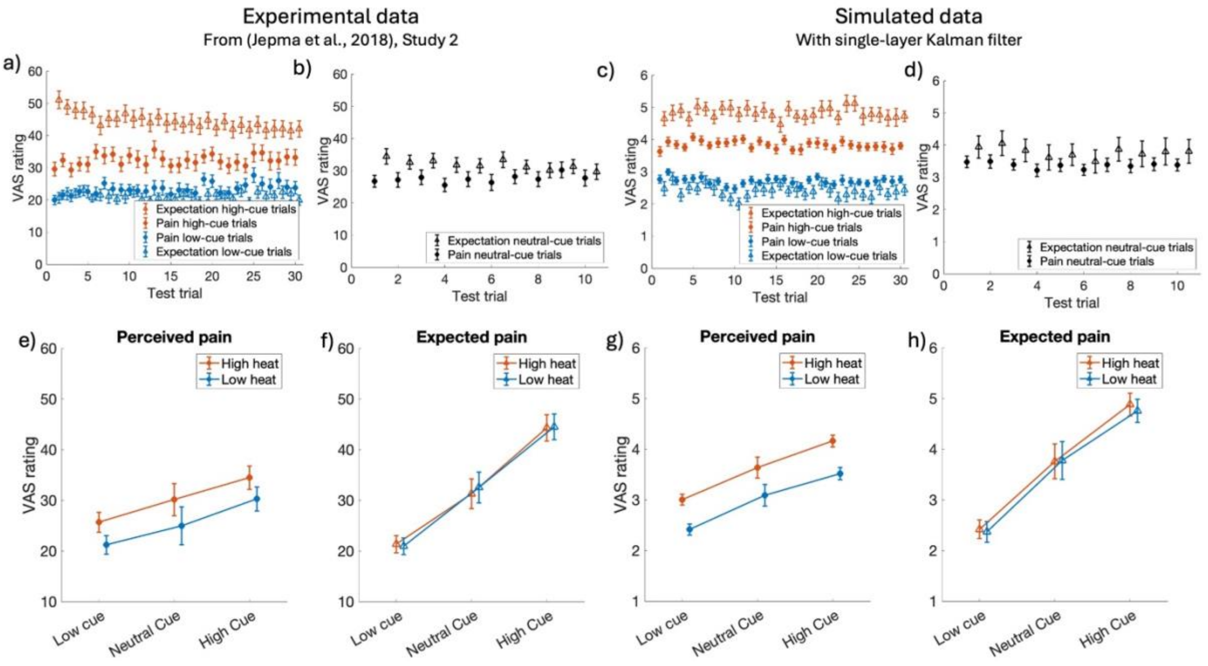


*Figure S 1 Experimental data from Jepma et al., Study 2* [13] (left) *and our Kalman filter simulations results (right).* ***a****),* ***b****),* ***c****) and* ***d****) average expected (open triangles) and perceived (filled circles) pain as a function of cue type and trial for experimental and simulated data, respectively.* ***e****) and* ***g****) perceived (filled circles),* ***f****) and* ***h****) expected (open triangles) pain as a function of stimulus temperature and cue type for experimental and simulated data, respectively. Error bars indicate inter-individual standard errors. Note that the experimental data is measured on a 100-unit scale, whereas the simulated data is on an 11-unit scale.*

### *Figure S2 – Study 2 from (Jepma et al., 2018) with hierarchical Kalman filter*

These simulations are run as described for Figure 4 in the main manuscript, but with the same modifications as for Figure S 1.

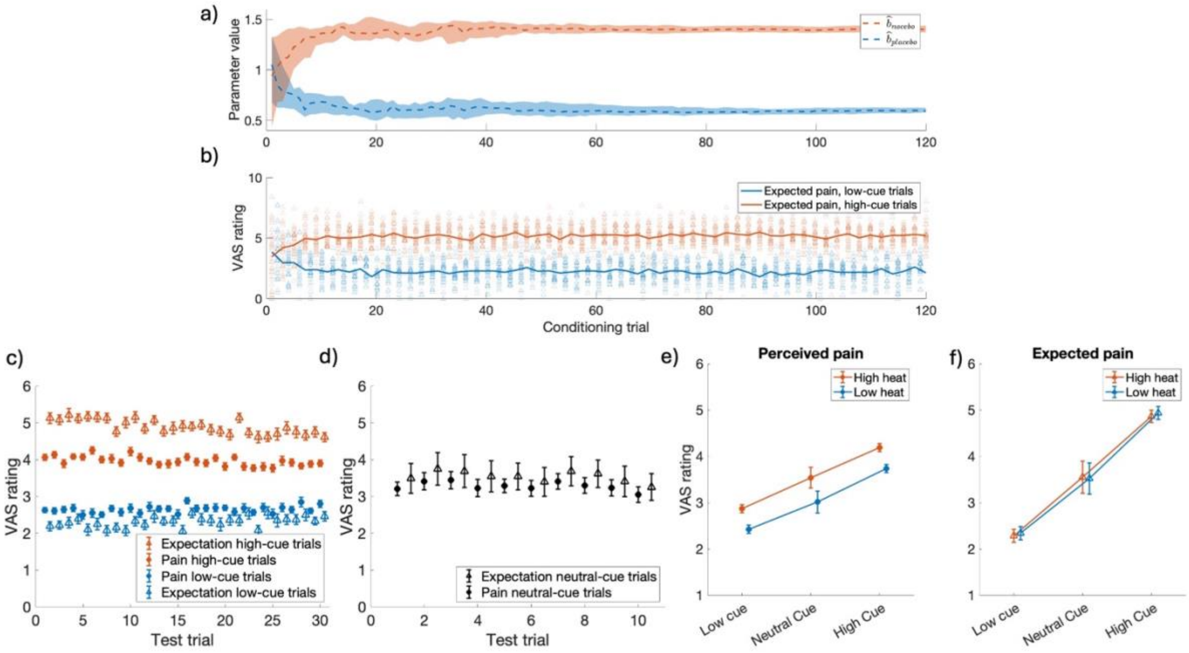


*Figure S 2 Results of the hierarchical Kalman filter simulations of classical conditioning, when using the data from (Jepma et al., 2018) Study 2 [13]. In the testing phase of Study 2 an additional, neutral cue is introduced, which was not included during the learning phase of the experiment.* ***a****) median value of $\hat{b}_{nocebo}$ (dashed red line) and $\hat{b}_{placebo}$ (dashed blue line) across conditioning trials. Shaded areas indicate the interquartile range.* ***b****) average expected pain for high-cue trials (red) and low-cue trials (blue) during conditioning. Open triangles indicate the expected pain for each participant on each conditioning trial.* ***c****) and* ***d****) the average expected ($\hat{\hat{x}}$, open triangles) and perceived ($\hat{x}$, filled circles) pain as a function of cue type on each test trial.* ***e****) perceived (filled circles,) and* ***f****) expected (open triangles) pain as a function of stimulus temperature and cue type. Error bars indicate inter-individual standard errors.*

### *Figure S3 – Chronic pain without control input*

Here, we run simulations just as for Figure 2 in the main manuscript, but with $u^{(k)} = 0 \; \forall \; k$. As described in the Methods section, this scenario reflects a situation where there are no predictive cues of an upcoming noxious stimuli $\tilde{u}^{(k)}$.

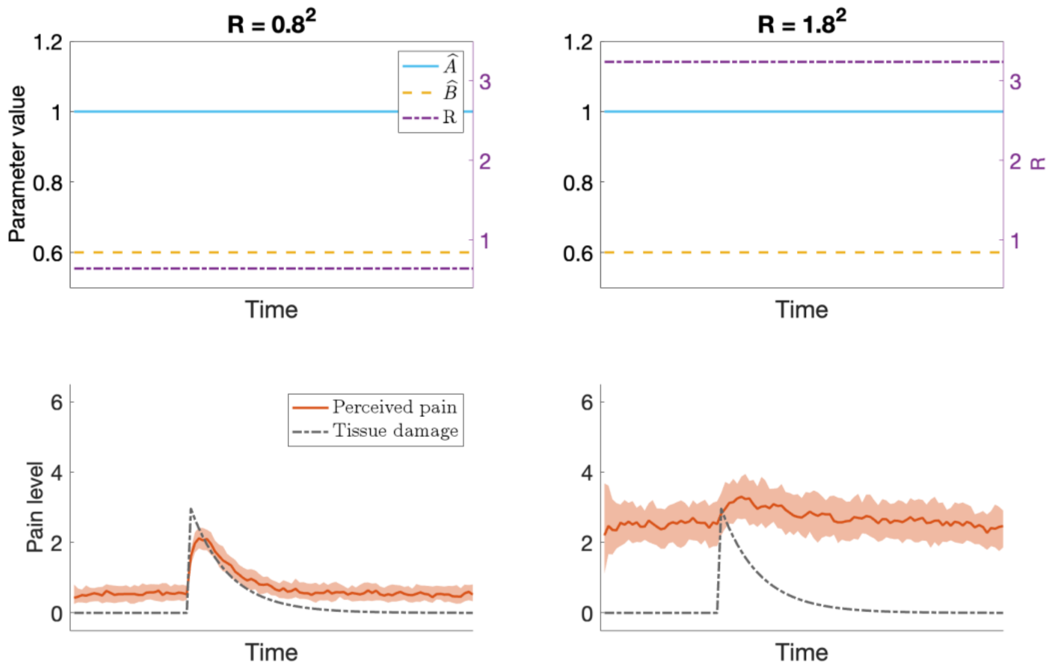


*Figure S 3 Results of the Kalman filter simulations of chronic pain with control input $u^{(k)} = 0 \ \forall \ k$. The results are similar to those presented in Figure 2 in the main manuscript. Despite having no predictive cues $u^{(k)} \neq 0$ of the upcoming noxious stimuli, the increase in tissue damage still gives rise to a sensory response that is sufficient to increase the level of pain. Additionally, when there is elevated uncertainty in the sensory input and the internal model parameter is $\hat{A} \approx 1$, the pain may persist even after the tissue damage has recovered.*
